# Competitive Microarray Screening Reveals Functional Ligands for the DHX15 RNA G-quadruplex

**DOI:** 10.1101/2023.07.25.550542

**Authors:** Peri R. Prestwood, Mo Yang, Grace V. Lewis, Sumirtha Balaratnam, Kamyar Yazdani, John S. Schneekloth

## Abstract

RNAs are increasingly considered valuable therapeutic targets, and in turn the development of methods to identify and validate both RNA targets and RNA-binding compounds is more important than ever. In this study, we utilized a bioinformatic approach to identify a hairpin-containing RNA G-quadruplex (rG4) in the 5^′^ UTR of *DHX15* mRNA. By using a competitive small molecule microarray (SMM) approach, we identified a compound that specifically binds to the *DHX15* rG4 with a K_D_ of 12.6 ± 1 µM. This rG4 directly impacts translation of a *DHX15* reporter mRNA *in vitro*, and binding of our compound (F1) to the structure inhibits translation up to 57% with an IC_50_ of 22.9 ± 3.8 µM. The DHX15 protein is an “undruggable” helicase associated with several types of cancer progression, and our data represent the first published effort to target the rG4 in *DHX15* mRNA to inhibit its translation. Overall, our work is informative for the development of novel small molecule cancer therapeutics for RNA targets starting from target identification.

**Graphical Abstract:** 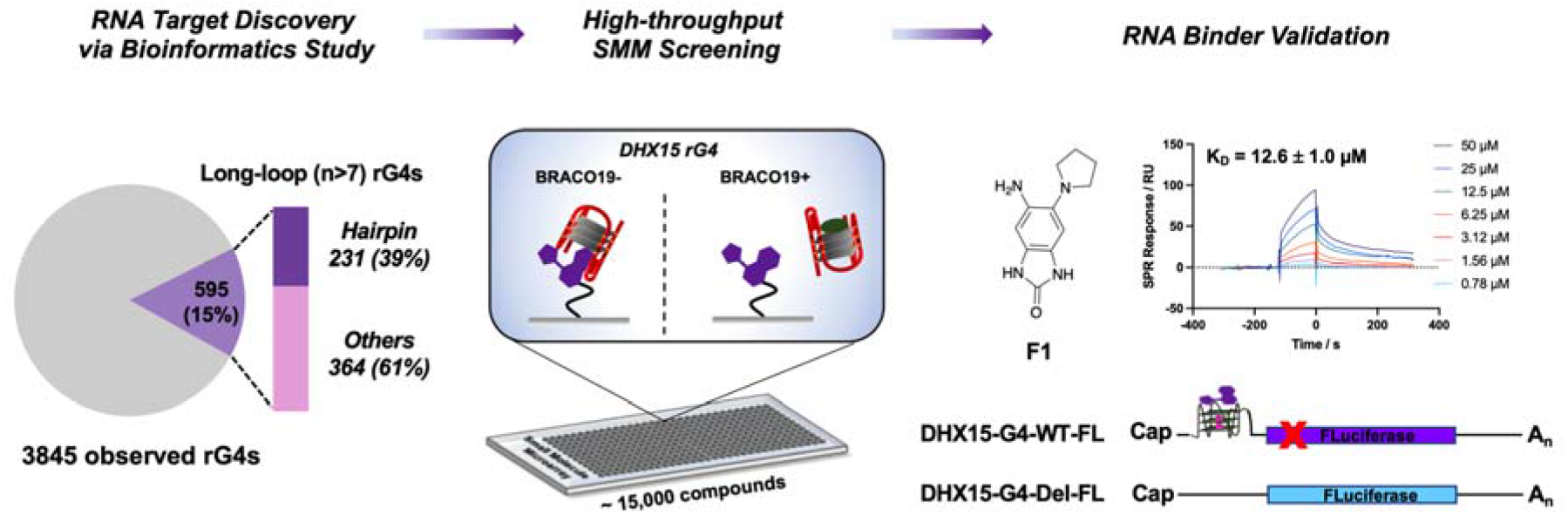

## 1. Introduction

In recent years, RNAs have become attractive targets for small molecule medicinal chemistry campaigns.^1^ This is partly due to the sheer amount of RNA in cells—approximately 85% of the genome is transcribed into RNA, compared to the approximately 1-2% of the genome that encodes proteins.^2, 3^ Further, just ∼15% of the proteome is “druggable,” leaving many disease-relevant proteins outside the realm of targetability with small molecules.^4^ Recent research suggests that several types of RNA (such as lncRNA, miRNA, and mRNA) can be effectively targeted with druglike molecules.^5, 6^ Targeting mRNA presents a powerful opportunity to control the expression of pathogenic, but “undruggable”, proteins.

Ideal RNA targets for small molecules should meet the requirements of both ligandability and functional relevance. Like proteins, many RNAs can fold into diverse secondary and three-dimensional structures with hydrophobic pockets in many structures throughout the protein databank (PDB).^6-8^ Moreover, these structures or conformations can be recognized by proteins or small molecules, and regulate diverse functions such as translation, splicing, and RNA decay.^9-13^ In some cases, targeting ligandable structures with small molecules can result in functional changes caused by altering the thermodynamic stability or conformation of the RNA.^14, 15^ For example, RNA G-quadruplexes (rG4s), which contain guanine-rich tracks that fold to form the high-order structure, have been regarded as attractive RNA targets. Several rG4s in 5’UTR regions has been targeted with small molecules (including the FDA-approved drug Levofloxacin), resulting in translation inhibition.^16-18^ In particular, rG4s with duplex/hairpin structures embedded in loops have also been attracting attention for their unique structures and protein binding (e.g., sc1 RNA-FMRP RGG, HIV-1 RNA U3).^19, 20^ These intriguing RNA structures exhibit great potential for small molecule targeting.

Once an RNA target is identified, high-throughput screening (HTS) is an effective way to identify potential hit compounds. Multiple biochemical and biophysical techniques have been reported to screen compound libraries against RNA targets.^21^ Our lab has published extensively on the use of small molecule microarray (SMM) technology to identify RNA-binding compounds in an efficient manner.^22, 23^ To discover hits, fluorescently labeled nucleic acids are incubated on glass slides that contain tens of thousands of small molecules spatially arrayed and covalently linked to the surface. Imaging and analysis of the slide reveals hit compounds bound to RNA. The development of SMM, new methodologies (e.g., muti-color imaging), and a large dataset of orthogonal RNA-small molecule binding has resulted in a more comprehensive understanding of RNA targeting.^23, 24^

In this study, we employed a bioinformatic approach to identify non-canonical RNA G-quadruplexes (rG4s) in the human transcriptome, revealing a hairpin-containing rG4 in the 5^′^ UTR of the mRNA for *DHX15*. We used multiple biochemical and biophysical techniques to establish the presence of this G4 in the *DHX15* mRNA and characterize its folding. To discover small molecule ligands for the *DHX15* rG4 with defined binding modes, we used a novel, biochemically competitive SMM screening approach. The most active identified compound (**F1**) bound reversibly to the rG4 (K_D_ = 12.6 ± 1 µM) and inhibited translation of a *DHX15* luciferase reporter *in vitro*, making it a promising hit. More broadly, our pipeline combining bioinformatics with biochemically competitive HTS efforts demonstrates a practical strategy for identifying targetable, structured regions within complex mRNAs and ligands with defined binding modes. Overall, our work is informative for the development of novel anticancer compounds from target identification to functional assays.

## 2. Materials and Methods

### 2.1. Materials

*DHX15* rG4 RNA (5’-CGU GUG GGC UGU AGU AGC GGG AGG GGU GGG G-3’) samples were purchased from Dharmacon with HPLC purification. RNAs were chemically synthesized with Cy5 or biotin labeling according to the requirements for high-throughput SMM screening and biomolecular binding assays, respectively. To prevent RNA degradation, all buffers used in this study were prepared with RNase-free water.

### 2.2. Bioinformatic study of targetable structures

Browser Extensible Data (BED) files were downloaded from the Gene Expression Omnibus (GSE77282) pertaining to the experimental rG4-seq data. The genomic sequences of the BED files were extracted from GRCh37 (hg19) reference genome using bedtools getfasta (bedtools 2.30.0) according to the original database. The folding test on long loops was based on previously reported methods.^25, 26^ G4-forming sequences were identified from the extracted rG4s using the regular expression formula (G_3-6_ N_1-33_)_3_ G_3-6_ (N stands for A, C, or U) and the potential for hairpin-forming loops within the rG4 sequences were analyzed. UNAFold 4.0 was used to calculate duplex formation from sequences flanked between two G-tracts. Loop sequences with more than 50% of their nucleotides base paired were identified as duplex-forming sequences. Then, the hairpin-rG4 sequences were traced back to the corresponding transcripts and genes. The high throughput folding test was performed on the NIH HPC Biowulf cluster. Coordinates of the hairpin-containing rG4s were then converted to GRCh38 (hg38) coordinates using the UCSC Lift Genome Annotations tool (liftOver).

### 2.3. Circular dichroism (CD) assay

For characterization of folded or unfolded RNA structures, a CD spectrometer (JASCO, USA) was used. In brief, unlabeled *DHX15* RNA was folded under a designated buffer condition by heating at 90^□^ followed by snap cooling on ice. The annealed RNA was kept at room temperature for 10 min before further experiments. For CD spectrum collection, each sample was scanned in a range of 230–320 nm in triplicate, followed by referencing the buffer spectrum. To test the effect of the compound on RNA conformational change, the RNA sample was mixed with the compound at different concentrations (keeping 5 µM RNA and 5% DMSO in the final solution).

### 2.4. High-throughput SMM screening

For high-throughput SMM screening, the microarray slides were prepared based on previously reported methods using a robotic microarray printer (Arrayjet, UK).^27^ On each slide with 2D-epoxy modification, over 7,000 compounds were printed in duplicate spots, forming a high-density microarray. The *DHX15* RNA target with a Cy5-label at the 5^′^ end was annealed prior to the screening. The SMM slide was first blocked with 500 nM yeast total tRNA (Invitrogen) in screening buffer (10 mM Tris, pH 7.0, 100 mM KCl, 0.005% Tween 20) for 2 h at room temperature. Then, the working solution was changed to Cy5-labeled *DHX15* RNA solution at 50 nM, followed by incubation for another 2 h. After that, the SMM slide was gently washed with PBST, PBS, and ddH_2_O three times each, then dried by centrifugation (1,700 g, 2 min). Finally, the microarray was imaged with an InnoScan 1100 AL fluorescence scanner (Innopsys, France) at 635 nm and the hits were identified based on the quantified fluorescent intensity of each spot. For competitive screening, Cy5-*DHX15* RNA was incubated with or without 500 nM BRACO19. After data acquisition, Z-scores from two individual screenings were plotted and compared. Competitive or non-competitive hits were identified using the SMM data analysis pipeline developed by the Schneekloth lab at NCI.

### 2.5. Surface plasmon resonance (SPR) assay

To determine the binding performance of the hit compounds against *DHX15* RNA, SPR experiments were carried out using a BIAcore 3000 instrument (Cytiva, USA). To avoid RNA degradation during long-lasting experiments, the whole microfluidic system was rinsed by priming 50% RNase Zap buffer and RNase-free water before starting. Then, the running buffer (10 mM Tris, pH 7.0, 100 mM KCl, 0.005% Tween 20, 5% DMSO) was primed into the system, keeping the flowrate at 5 µL/min. Next, a CM5 chip containing two flow-cells (Fc1 and 2) was activated (0.4 M EDC and 0.1 M NHS) for 10 min and then coupled with streptavidin (SA) solution (200 µg/mL in 10 mM sodium acetate buffer, pH 4.5) for another 20 min, resulting in ∼6000 response units (RUs) signal. After SA immobilization, the chip surface was chemically quenched (1 M ethanolamine, pH 8.5) for 10 min and regenerated (10 mM NaOH) for 2 min. Then, biotin labeled *DHX15* RNA was annealed at 5 µM and only injected into Fc 2 for 30 min while Fc1 was left blank as a reference channel. The RNA immobilization could reach ∼2000 RU on the chip surface. The system continued flowing the running buffer until the baseline was stable prior to future steps.

To detect binding between small molecules and *DHX15* RNA, a higher flowrate of 25 µL/min was used to avoid mass transfer effect. The compound solution (50 µL) was applied to both Fc1 and Fc2 for 120 s association, followed by 200 s dissociation with buffer injection. In addition, 1 M KCl solution was used (pulse injection) to regenerate the surface if needed. To measure the dissociation constant (K_D_), compound solutions with different concentrations were injected successively, and the K_D_ value was calculated by fitting the binding curves based on Langmuir 1:1 binding model in BIAevaluation software (Cytiva, USA).

### 2.6. Fluorescent intercalator displacement (FID) assay

In this study, an FID assay was used to validate the binding mode/site of RNA-small molecule interactions. Herein, thioflavin T (ThT) was used as a fluorescent probe that can bind to the G-quartet of *DHX15* RNA. To test the displacing capability of the compounds, ThT and pre-folded *DHX15* RNA were mixed in the running buffer, resulting in 500 nM and 250 nM, respectively. Small molecules from SMM screening were prepared in a black 96-well plate (Costar, USA) at a designated concentration, followed by addition of the RNA-dye solution, resulting in a total volume of 100 µL in each well. The plate was kept on a shaker for 15 min at room temperature. After a quick centrifugation (1,700 g, 2 min), the plate was loaded into a Synergy Mx microplate reader (BioTek, USA). The fluorescent intensity for each well was recorded at Ex 430 nm/Em 500 nm.

### 2.7. Plasmid construction

To construct the plasmid p^DHX15-G4-FL^, which encodes the 161-nucleotide UTRQ transcript, we received the 185 bp g-block from IDT which contain 161 bp of *DHX15* 5^′^ UTR with *HindIII* and *BamH1* tailed at 5^′^ and 3^′^ respectively. The g-block was amplified by two specific primers: forward primer (5^′^-ATATATAAGCTTCTCGTCGCCGCCG-3^′^) and reverse primer (5^′^-ATATATGGATCCCCTCGCACTCTTCGAAC-3^′^) and purified by gel electrophoresis. This amplified product was digested with *HindIII* and *BamH1* and purified by gel electrophoresis. In ligation, we ligated the gel-purified fragments at the *HindIII* and *BamH1* site and inserted into the *HindIII* and *BamH1* digested linear p^T7-CMVtrans-FFLuc-polyA^ vector (Addgene 101156). We confirmed positive clones for the full length of the insert by DNA sequencing. To construct p^DHX15-G4-Del-FL^ the control plasmid encoding the transcript DHX15-G4-Del-FL, we deleted the 31 base pairs of the *DHX15* 5^′^ UTR, including the G-quadruplex-forming sequence. The 154 bp g-block from IDT which contain 130 bp of *DHX15* 5^′^ UTR with *HindIII* and *BamH1* tailed at 5^′^ and 3^′^ respectively. The g-block was amplified by specific primers and ligated into the linear p^T7-CMVtrans-FFLuc-polyA^ vector as described above. We sequenced all the constructs to confirm the presence of the intended changes.

### 2.8. In vitro transcription

The plasmids p^DHX15-G4-FL^ and p^DHX15-G4-Del-FL^, which encode the transcripts DHX15-G4-FL and DHX15-G4-Del-FL, respectively were constructed as described above. The plasmids were then linearized at the 3’ end using *Btgz*, and linearized plasmids were used as a template for *in vitro* transcription. The 5’-capped transcripts were synthesized *in vitro* using mMessage mMachine T7 (ThermoFisher Scientific), followed by incubation with DNase for 15 min at 37°C to remove the template DNA. Then, all the transcripts were purified using Monarch RNA Cleanup Kit (NEB). The concentration was determined using a Nanodrop. Integrity and size of each transcript was confirmed using a 1% agarose gel.

### 2.9. In vitro translation and luciferase assay

*In vitro* translation of DHX15-G4-FL and DHX15-G4-Del-FL transcripts in the presence of **F1** or BRACO19 was carried out in a cell-free translation system of rabbit reticulocyte lysates (Promega) following manufacturer’s protocol. Briefly, 500 ng of DHX15-G4-FL and DHX15-G4-Del-FL transcripts were folded by heating at 90ºC for 3 min then slowly cooling down to room temperature for 1 h. Next, **F1** (2 µM to 100 µM) or BRACO19 (10 µM) was added into the folded RNA and incubated for 15 min at room temperature. Samples incubated with 5% DMSO were used as a control. The *in vitro* translation was carried out by adding the reticulocyte lysate into the samples followed by incubation at 30ºC for 90 min. The firefly luciferase activity was measured using luciferase assay reagent (Promega) according to manufacturer’s protocol on a Synergy Mx microplate reader (BioTek).

## 3. Results and Discussion

### 3.1. Discovery of an rG4 structure in DHX15 mRNA

Many eukaryotic RNAs have been observed to contain rG4-forming sequences.^28, 29^ To identify novel rG4s as targets, we assessed a public rG4-seq dataset assembled to identify rG4s in human mRNAs.^30^ As shown in **Fig. 1A**, a total of 3,845 rG4s were observed to form in K^+^ buffer by a transcriptional stalling method. While most rG4s were canonical quadruplex structures (G_3-5_N_1-7_G_3-5_N_1-7_G_3-5_N_1-7_), about 15% (595) of the rG4s were found to contain longer loops (n>7). For those long-loop rG4s, 39% had the potential to form duplex (or hairpin) structures by base pairing within loops, making their 3D structures distinct from simpler G4s. Among the 230 putative hairpin-containing rG4 sequences corresponding to 72 genes (**Supplementary File 1**), a short RNA sequence from DEAH-box helicase 15 (*DHX15*) mRNA was notable for three reasons. First, DHX15 protein is an RNA helicase involved in mRNA processing and immunity.^31, 32^ It has been reported that DHX15 is overexpressed in and leads to the progression of several cancer types, including prostate cancer, Burkitt lymphoma, and acute myeloid leukemia.^33-36^ Second, as an RNA helicase the DHX15 protein is difficult to target directly with small molecules.^37^ Finally, the rG4 is embedded within the 5^′^ untranslated region (5^′^ UTR) of *DHX15* mRNA (**Fig. 1B**), which could potentially regulate its translation, as is the case with other rG4s.^38^ In addition, this quadruplex-forming sequence was found in 17 different cell lines by RT-PCR experiments (**Supplementary Fig. 1**). Together, these factors indicated that the *DHX15* rG4 could be a good target candidate for small molecules.

**Fig. 1.**
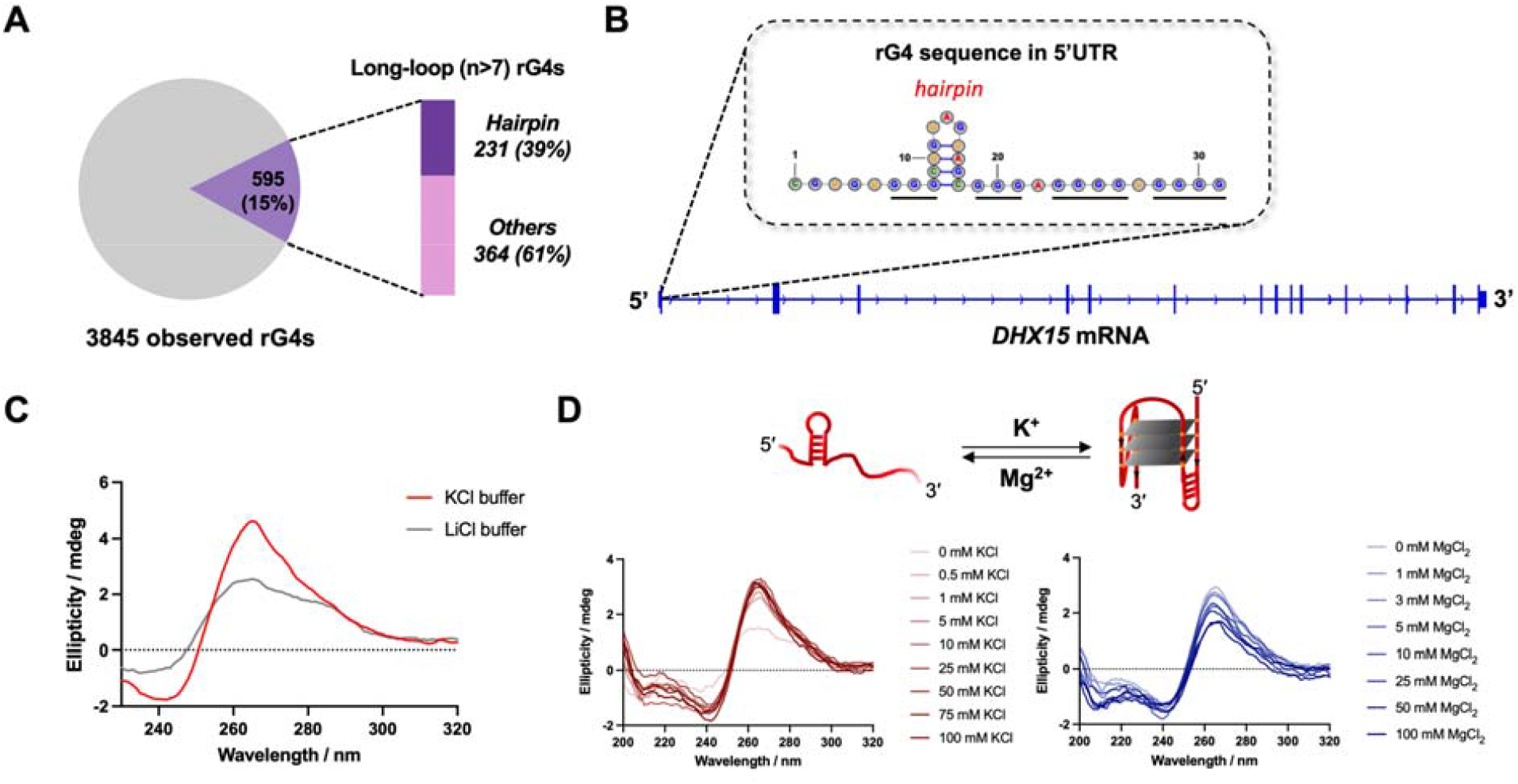
Identification of an rG4 in *DHX15* mRNA and *in vitro* study of G4 formation. (**A**) Bioinformatic study on the rG4s with long loops formed in human cells. The observed rG4s were obtained from reported rG4-seq data. (**B**) The hairpin-containing rG4 (secondary structure) discovered in the 5’ UTR region of *DHX15* mRNA. (**C**) CD spectrum characterization of the folded/unfolded *DHX15* rG4 RNA in KCl or LiCl buffer. (**D**) Cationic ion titration assay using K^+^ and Mg^2+^ to induce the RNA confirmational change between stem-loop and hairpin-rG4 structures.

Characterization of *DHX15* rG4 folding was performed using circular dichroism (CD) spectroscopy. To assess the folding conditions of the RNA, a short construct containing the *DHX15* rG4 was annealed and folded in buffer containing either KCl or LiCl prior to recording the CD spectrum. As shown in **Fig. 1C**, RNA folded in KCl buffer exhibited a characteristic G4 structure, demonstrated by a positive peak at 263 nm and a negative peak at 240 nm. In contrast, a broadened peak was observed in the spectrum corresponding to the RNA folded in LiCl buffer, implying the presence of a less folded RNA (**Fig. 1C**). A conformational shift between quadruplex and duplex structures was also identified by utilizing a cationic ion titration assay demonstrating that *DHX15* rG4 formation is induced by the addition of K^+^ and inhibited by Mg^2+^ (**Fig. 1D**).

### 3.2. High-throughput screen identifies small molecule ligands for the DHX15 rG4

Having demonstrated folding of the rG4 *in vitro*, a high throughput SMM screen was performed to identify small molecule ligands (**Fig. 2A**). SMM slides were fabricated as previously described,^22^ and a total of 14,802 compounds were screened. In one condition, slides were incubated with Cy5-*DHX15* rG4. In parallel, slides were incubated with both Cy5-*DHX15* rG4 and BRACO19, a well-studied, pan-G4 ligand that stacks on tetrads. After incubation, scanning and data analysis revealed hits by quantification of fluorescence intensities of the spots on the slides. Z-scores were plotted on a 2D scatter plot, wherein the points in the highlighted area represent small molecules that bound to the same site as BRACO19 as evidenced by their lower Z score in the presence of the competing ligand (**Fig. 2B**). Among the screened compounds, 195 of them (hit rate: 1.3%) showed competitive behavior against BRACO19 (**Supplementary File 2**). In addition, binding selectivity for each hit was determined by comparing the Z-scores from the *DHX15* rG4 screen with the Z-scores from a compiled dataset of 23 other nucleic acid screens.^24^ By analyzing spot morphology and structural diversity, we recognized 17 compounds as competitive hits, yielding a final hit rate of 0.1%. Single-concentration screening by SPR and a fluorescence quenching assay were performed on the hits to evaluate their binding ability towards the *DHX15* rG4 (**Supplementary Figs. 2 and 3**). Following these assays, we focused our study on compound **11**, which demonstrated good competitive binding performance (Z-score_rG4_ = 3.9, Z-score_rG4+BRACO19_ = 1.3) and excellent selectivity (Gini = 0.939) in the high-throughput screen **(Figs. 1C and D**).^24, 39^

**Fig. 2.**
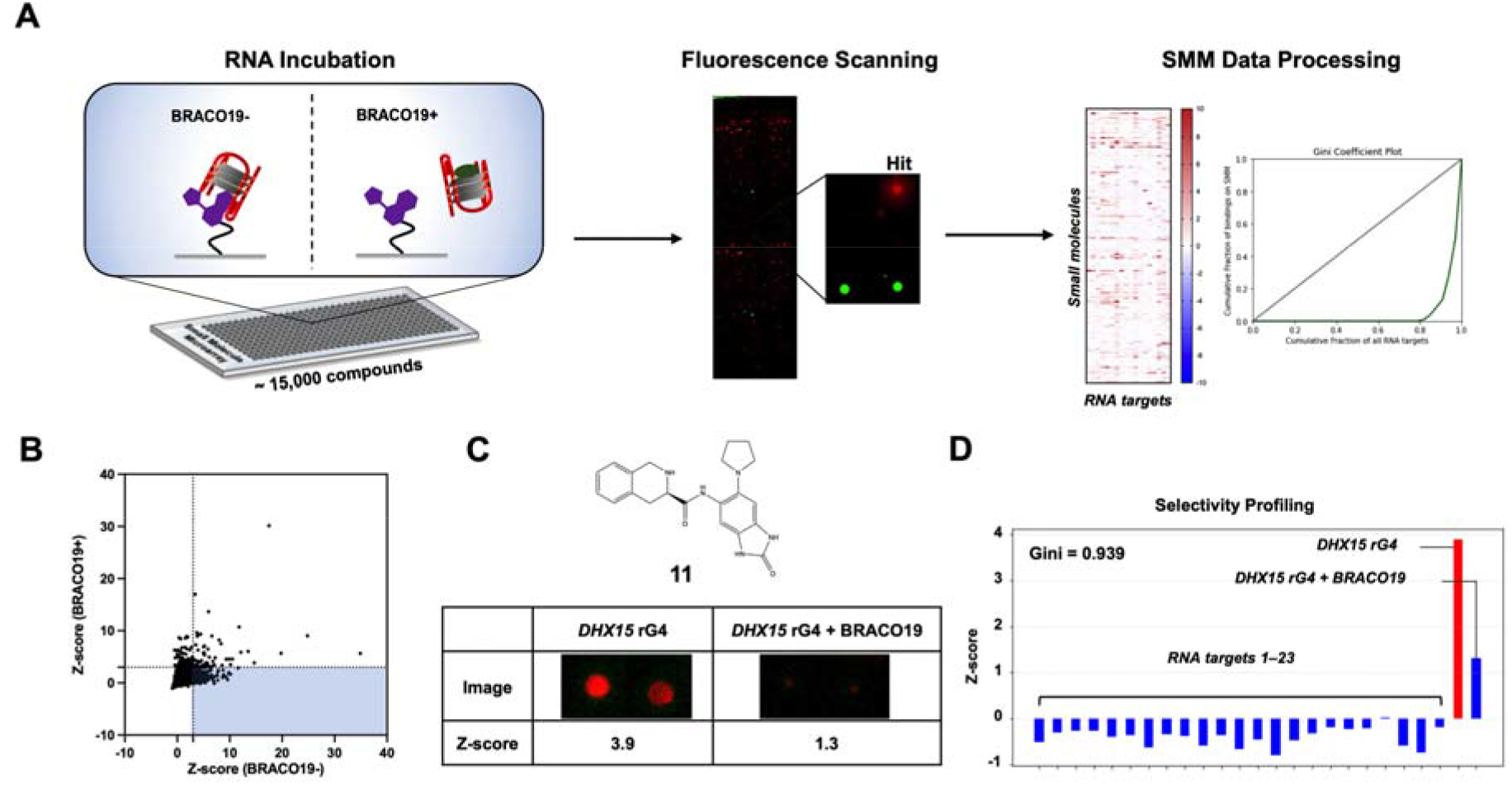
High-throughput screening using SMMs to identify *DHX15* rG4 binders. (**A**) Schematic representation of SMM workflow for rG4 target screening. A competitive screening strategy was used by incubating the RNA in the presence/absence of BRACO19. (**B**) Scatter plot of screening results showing the comparison of Z-scores in presence/absence of BRACO19. (**C**) Identification of compound **11** as a hit from SMM screening. (**D**) Selectivity profiling of compound **11** based on the compiled screening dataset.

### 3.3 Biophysical analysis of ligand binding to the DHX15 rG4

To confirm the validity of the original hit, compound **11** was resynthesized to test in binding assays. Compound **11** was synthesized by first performing an amide coupling between a Boc-protected amino acid and the aniline **F1** shown in **Fig. 3A**. After deprotection and reverse phase purification, compound **11** was isolated with a 34% yield over two steps (**Fig. 3A**). Upon preparation of stock solutions, it was observed that compound **11** readily decomposed in DMSO at room temperature upon standing. Liquid chromatography/mass spectrometry (LC/MS) revealed decomposition in solid and solution stocks of both the resynthesized and commercially available versions of compound **11** (**Supplementary Fig. 4**). The rapid degradation of this compound, especially in solution, resulted in poor reaction yields (**Fig. 3A**) and an inability for us to reliably test the compound in biophysical or biological assays. Because the initial screen identified a competitive rG4 ligand with a good Z-score, we hypothesized that the original compound decomposed in such a way that a fragment of the compound remained immobilized to the slide and bound to the RNA during screening. Further, SPR assays were performed on pure compound **11** and a decomposed stock of compound **11**, a much higher binding signal was observed for the decomposed compound (**Supplementary Fig. 4**).

**Fig. 3.**
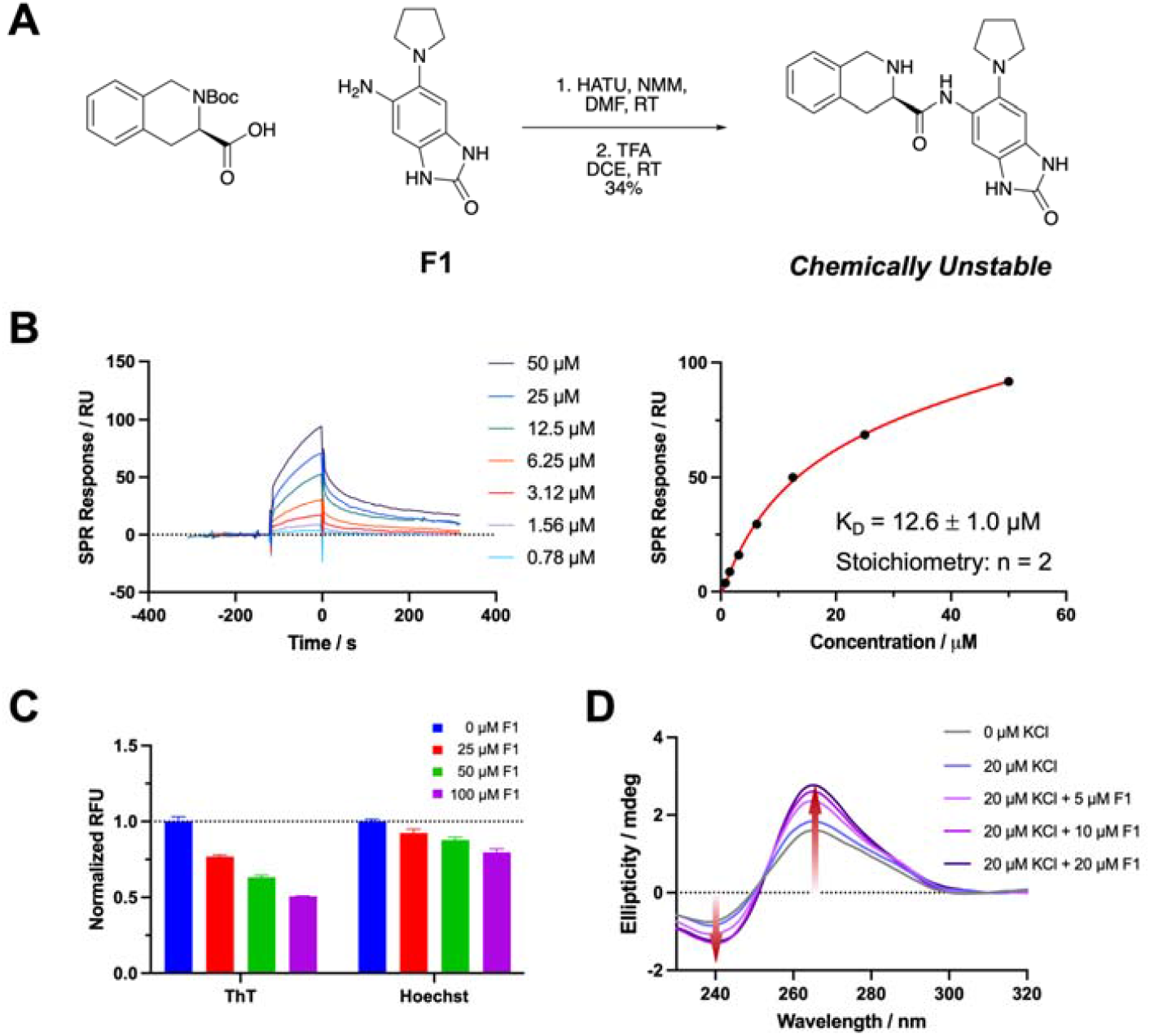
Identification of active compound via binding assays. (**A**) Synthetic preparation of compound **11**. (**B**) Titration of **F1** on SPR with dose-dependent binding curves and K_D_ determination by fitting at steady state. (**C**) Dose-dependent FID assay using two fluorescent probes (ThT and Hoechst 33258). (**D**) CD spectra showing the *DHX15* rG4 formation induced by addition of **F1** at increasing concentrations.

As a starting point for the identification of the active fragment, we performed SPR on the compounds used to synthesize compound **11**. The benzimidazol-2-one fragment (**F1**) and the *R* and *S* enantiomers of the deprotected tetrahydroisoquinoline (**F2** and **F3**) were each evaluated for their ability to bind to the *DHX15* rG4 (**Supplementary Fig. 5**). Here, **F1** displayed a strong binding signal and better binding kinetics than the other two fragments at 50 µM (**Supplementary Fig. 5**). By fitting the titration curves, **F1** bound with a 2:1 stoichiometric ratio and had a K_D_ of 12.6 ± 1 µM (**Fig. 3B**), while **F2** and **F3** had much lower binding affinities (K_D_ >100 µM) (**Supplementary Fig. 6**). The observed binding activity of **F1** to the rG4 was therefore deemed significant enough to continue our study of this fragment alone.

Once we determined that **F1** bound to the *DHX15* rG4 by SPR, we focused on identifying its mode of binding. During screening, the original hit was competed away by BRACO19, demonstrating that the hit was a G4 binder and likely stacked on the tetrads like BRACO19. To verify that **F1** forms a stacking interaction with the rG4, we performed an FID assay using ThT, a fluorescent G4-stacking compound. The results showed that ThT was displaced by up to 50% when **F1** was titrated into solutions containing ThT and *DHX15* rG4, indicating that **F1** also stacks on G4 tetrads (**Fig. 3C**). In contrast, only ∼20% displacement was observed when 100 µM **F1** was added to a solution of the rG4 and Hoechst 33258, a fluorescent groove-binding ligand. To further investigate the effect of **F1** on rG4 folding, CD spectroscopy was performed. **F1** was titrated into a solution of *DHX15* RNA in buffer containing 20 µM KCl, which meets the minimum requirement for rG4 folding. With increasing concentrations of **F1**, we observed the induction of rG4 formation, shown by the increased intensity of the characteristic spectrum (**Fig. 3D**). This result suggests that **F1** can shift the conformational equilibrium towards rG4 by binding and stabilizing its structure.

### 3.4 The rG4 from the DHX15 5’ UTR impacts translation in vitro

Because the rG4 is embedded in the 5^′^ UTR region of *DHX15*, we hypothesized that the structure might impact mRNA translation. To assess this, we designed two firefly luciferase reporter constructs—one containing the *DHX15* rG4 (DHX15-G4-WT-FL) and one lacking the rG4 (DHX15-G4-Del-FL) (**Fig. 4A**). Following plasmid construction and *in vitro* transcription, the resulting RNA constructs were tested in an *in vitro* translation assay. Each RNA was incubated with rabbit reticulocyte lysates, then firefly luciferase activity was measured for both constructs, with greater luciferase activity corresponding to greater translation. We found that compared to the wild-type construct, the rG4 deletion construct showed double the luciferase activity (**Fig. 4B**). This result suggests that the presence of the rG4 in the 5^′^ UTR of *DHX15* inhibits translation efficiency, making it a valuable small molecule target.

**Fig. 4.**
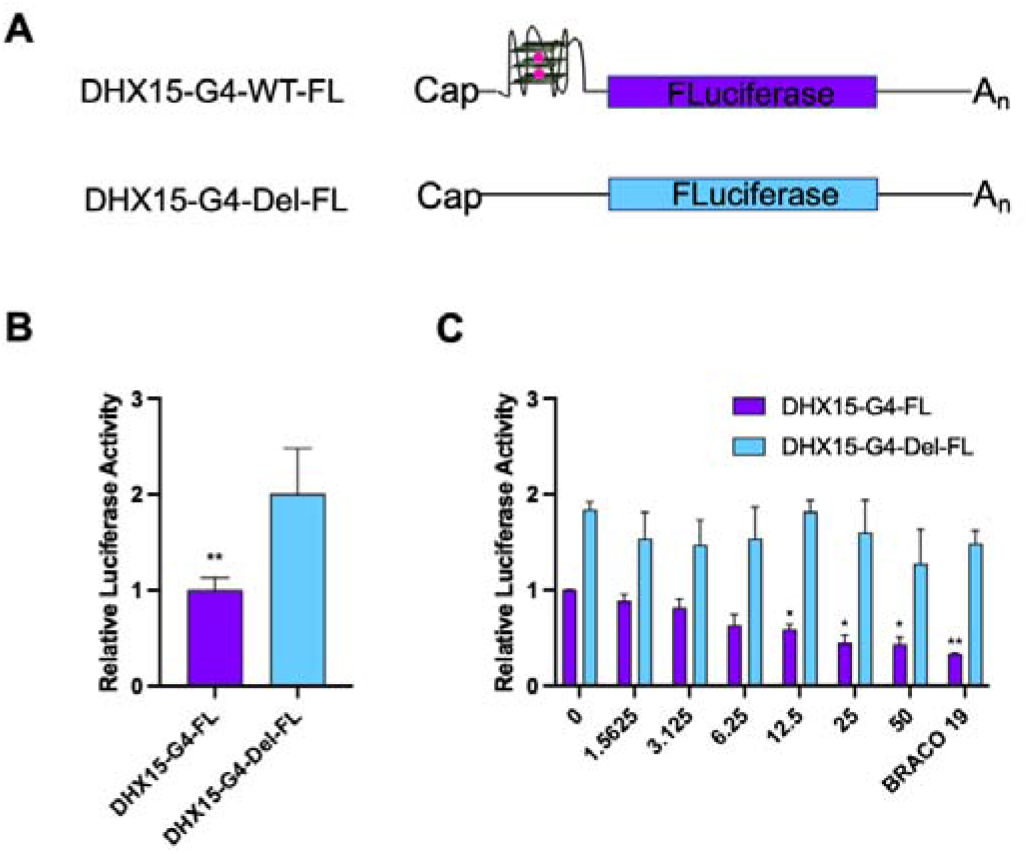
Translation inhibition assay using in vitro luciferase reporter system. (**A**) Schematic illustration of the *DHX15* luciferase reporter system RNA constructs. (**B**) Relative luciferase activity of WT and G4 deletion RNA constructs. (**C**) Dose-dependent testing of **F1** on translation inhibition of *DHX15* mRNA. *P < 0.005, **P < 0.0005.

To ascertain the effect of **F1** on *DHX15* mRNA translation, we applied the compound in the luciferase reporter system and performed a dose-dependent *in vitro* translation assay. As a result, the treatment with **F1** led to a dose-dependent response of translation inhibition (IC_50_ = 22.9 ± 3.8 µM), reaching a maximum inhibition value of 57% (**Fig. 4C, Supplementary Fig. 7**). In contrast, the translation efficiency of DHX15-G4-Del-FL was not significantly affected by treatment with **F1**. BRACO19 was used as a positive control to compare its mode of inhibition with that of **F1**. At 10 µM, BRACO19 inhibited translation of the rG4-containing construct by ∼67% but did not inhibit translation of the deletion construct (**Fig. 4C**). Based on these data, we confirmed that **F1** binds to the rG4 specifically and inhibits translation in a dose-dependent manner.

## 4. Conclusion

Here, we demonstrate a pipeline for the rapid discovery of rG4-targeting small molecules with defined binding modes by combining a bioinformatic approach, high-throughput SMM screening, and biophysical and functional assays. With this strategy, we identified a unique rG4 structure in the *DHX15* 5’ UTR as a cancer-relevant target and successfully discovered a compound that can downregulate the expression of DHX15 protein. To our best knowledge, no small molecule probes targeting *DHX15* mRNA have been reported.

Extensive work on optimizing the lead compound (**F1**) would be needed to achieve improved binding affinity, improve functional efficacy, and characterize specificity. Full utilization of the chemical space within the hairpin-quadruplex junction could be the key to making a better ligand, studies that might be informed by investigation of the potential non-competitive hits from the SMM screen. This fragment-based strategy has previously been proven successful for the development of high-affinity, selective RNA-binders.^40-42^ Additionally, the investigation on the mechanism of translation inhibition needs to be continued. The binding of **F1** to the rG4 could potentially affect protein-RNA interactions thus modulating the translation of *DHX15*, though this specific mechanism remains unclear.

Overall, the methodology outlined in this paper will be broadly useful as a strategy for future RNA target identification and drug discovery efforts. The use of bioinformatic approaches to explore *in vitro* and *in cellulo* chemical probing datasets can facilitate the discovery of other ligandable RNA structures beyond G4s, such as bulges, pseudoknots, and three-way junctions. Additionally, biochemically competitive high-throughput SMM screening results in the discovery of compounds that bind a specific RNA site with a distinct binding mode. Knowledge of the binding mode allows for the efficient planning of biophysical, biological, or functional assays that can accurately evaluate hits for various RNA structures. Altogether, this strategy presents a streamlined approach for the drug discovery process from target identification to lead compound evaluation.

## Supporting information

DHX15 manuscript SI File

## Acknowledgements

We thank the members of the biophysics resource (Dr Sergey G. Tarasov and Marzena Dyba) for helpful comments and suggestions on biophysical experiments. This work also utilized the computational resources of the NIH HPC Biowulf cluster (http://hpc.nih.gov).

## Supplementary materials

Supplementary materials associated with this article can be found, in the online version, at doi: xxx.

