## Supplementary material for "Competitive Microarray Screening Reveals Functional Ligands for the DHX15 RNA G-quadruplex": DHX15 manuscript SI File

**Supplementary Methods**

***RT-PCR***

All cell lines (MCF-7, PC-3, C4-2, DU 145, LNCap, SKE-MEL-2, SiHa, H5518T, DU142, T47D, Bt 549, HEK-293T, HEK-293, ADR, H226, UOK 262, LS174T and HT1080) were grown in T75 flasks with proper medium according to the ATCC protocol. Once the cells reached 80% confluence, total RNAs were isolated using TRIzol reagent (Thermo Fisher Scientific), and DNA was removed by TURBO DNase (Thermo Fisher Scientific) digestion according to the manufacturer's protocol. First strand cDNA was then synthesized by Superscript first-strand synthesis kit (Invitrogen) with oligodT primers according to the manufacturer’s protocol. Briefly, 1 µg of RNA and oligodT were mixed and heated at 65℃ for 5 min to denature the RNA. Then, samples were placed in ice for 10 min. In a separate tube, a 2X reaction mixture was prepared by adding 2 µL of 10X RT buffer (containing the LiCl), 4 µL of 25 mM MgCl_2_, 2 µL of 0.1 M DTT, and 1 µL of RNaseOUT. The resulting 9 µL of reaction mixture was added into the RNA/primer samples and incubated at 42℃ for 2 min. Then, 1 µL of SuperScript II RT was added into the samples and incubated at 42℃ for 50 min, followed by incubation at 70℃ for 15 min to heat inactivate the enzyme. Next, 2 µL of cDNA was used for PCR amplification with Platinum SuperFi II DNA polymerase (Invitrogen). The reaction was incubated at 98℃ for 30 s, followed by 25 cycles of 98℃ for 10 s, 63℃ for 35 s and 72℃ for 30 s. Finally, samples were loaded into the 2% agarose gel and run for 1 hour. The specific set of primers that were used to amplify the 5′ UTR and CDS of the *DHX15* mRNA and CDS of DHX15-G4-FL reporter construct mRNA are shown in Supplementary Table 1.

**Supplementary Tables**

**Supplementary Table. 1** Primers used for RT-PCR

| **Name** | **Sequence (5**′**′to 3**′**′)** |
| --- | --- |
| Primer A | ACCGTGTGGGCTGTAGTA |
| Primer B | TCTCGATCTCGATCCTTCCC |
| Primer C | CCGTTCACCAACTTACCCCA |
| Primer D | ACTCATTGCAGCCACTCTCC |

**Supplementary Table. 2** Compounds tested for binding validation.

| **Compound No.** | **ID** | **Chemical Structure** | **Z-score** |
| --- | --- | --- | --- |
| 1 | 5465521 | 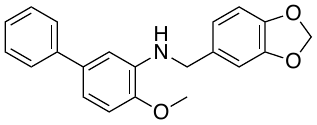 | 5.96 |
| 2 | 7134266 | 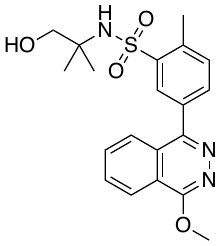 | 5.35 |
| 3 | 7657128 | 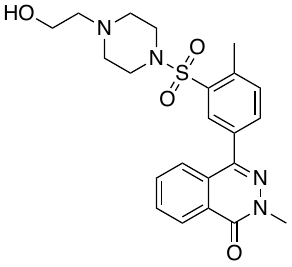 | 4.26 |
| 4 | 7997235 | 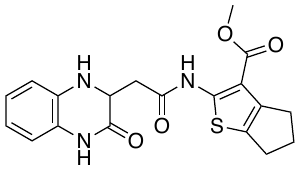 | 3.08 |
| 5 | 36223280 | 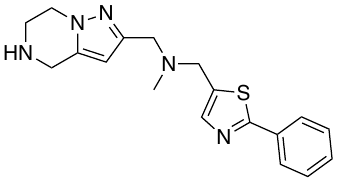 | 3.77 |
| 6 | 52156781 | 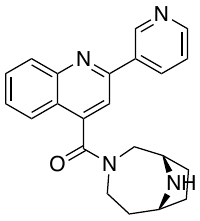 | 3.04 |
| 7 | 61724459 | 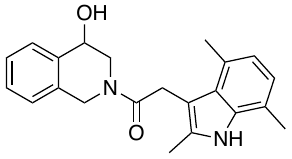 | 4.22 |
| 8 | 83545590 | 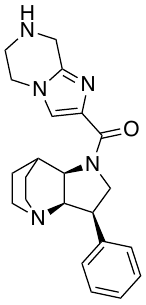 | 4.96 |
| 9 | 12317528 | 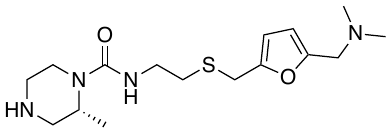 | 3.47 |
| 10 | 41464724 | 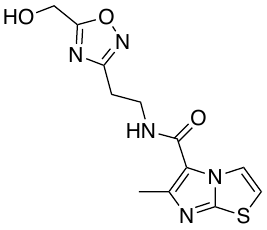 | 3.20 |
| 11 | 84731880 | 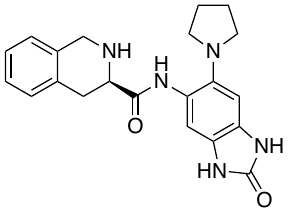 | 3.89 |
| 12 | 32052616 | 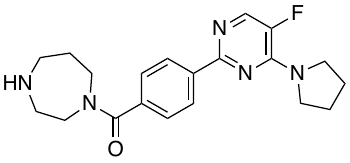 | 4.51 |
| 13 | 10287347 | 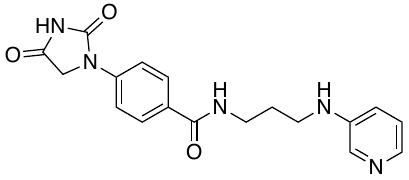 | 3.57 |
| 14 | 44321801 | 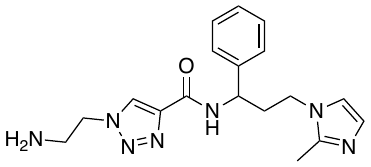 | 4.14 |
| 15 | 94198294 | 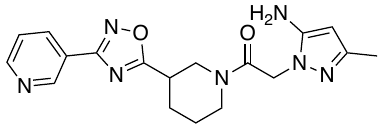 | 3.50 |
| 16 | 68270800 | 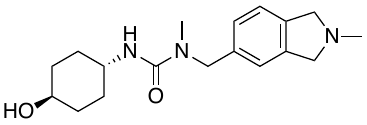 | 3.33 |
| 17 | 27813381 | 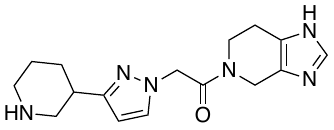 | 5.03 |

**Supplementary Figures**

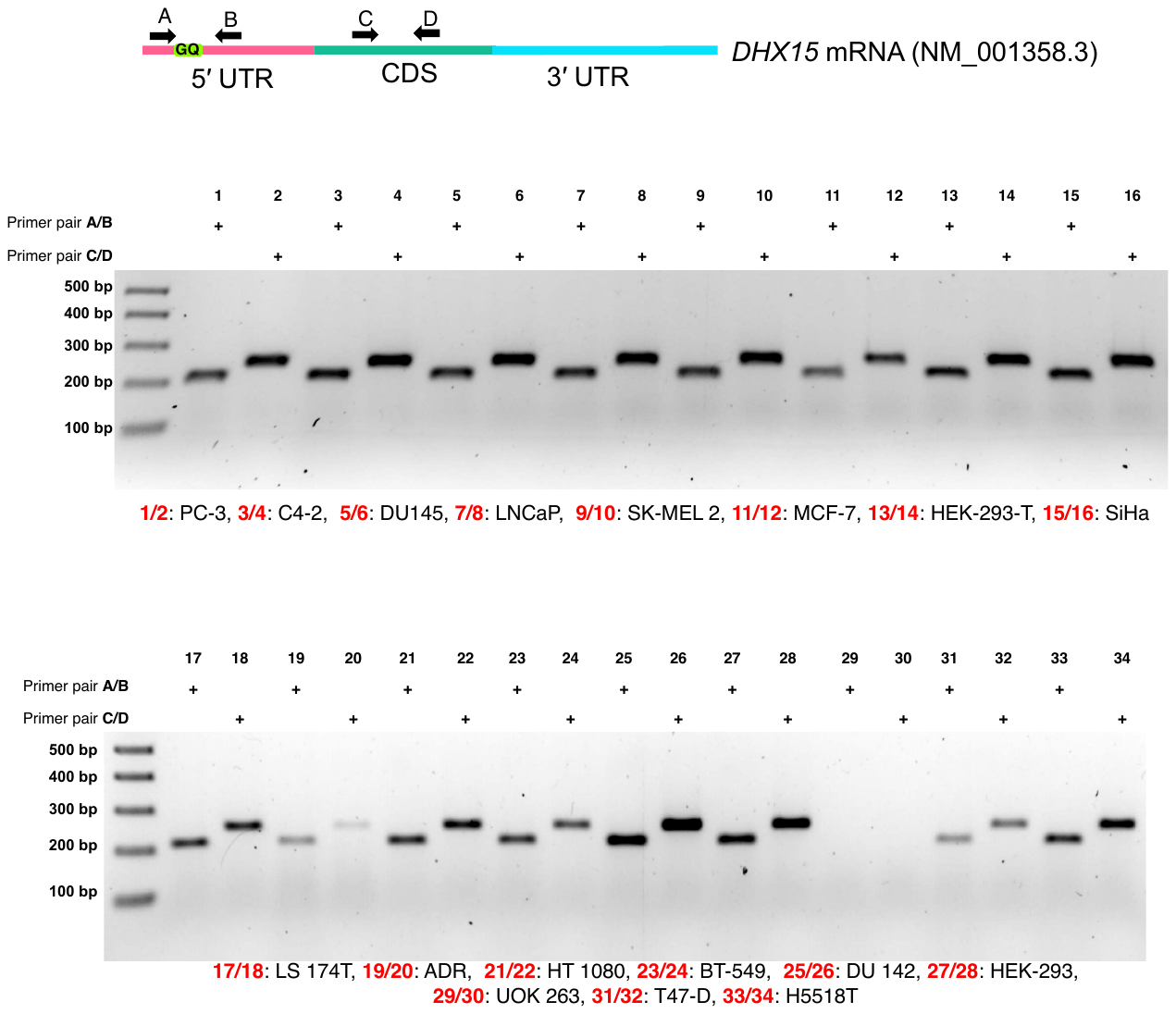

**Supplementary Fig. 1** RT-PCR results demonstrating the existence of the rG4-forming sequence in *DHX15* mRNA in 17 different cell lines.

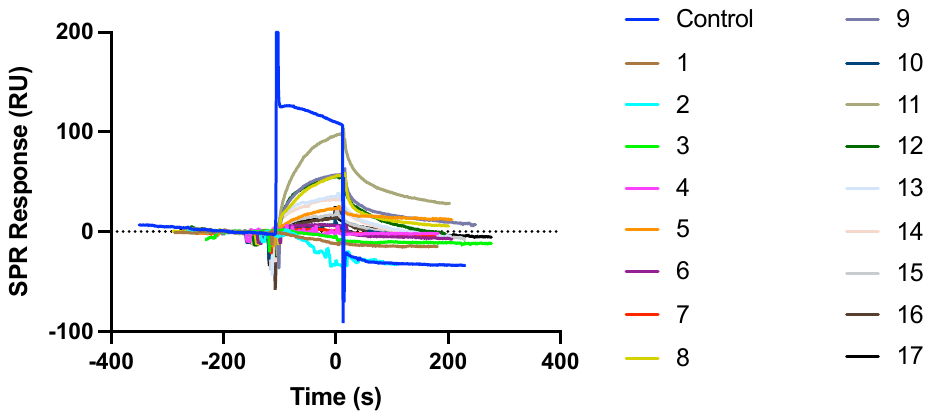

**Supplementary Fig. 2** Single-concentration (50 µM) screening by SPR to validate the binding between SMM hits and the *DHX15* rG4.

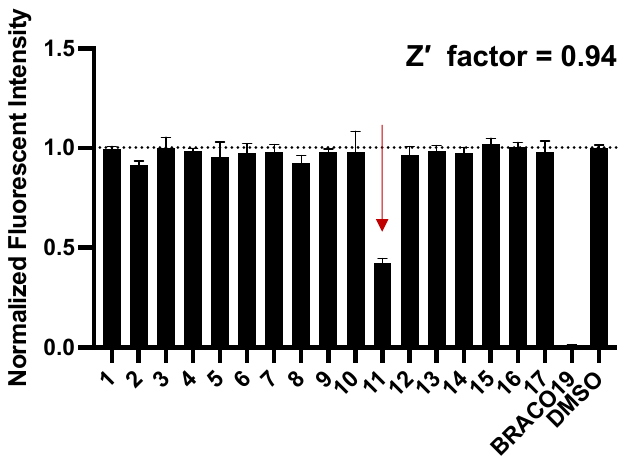

**Supplementary Fig. 3** Fluorescent intensity assay using 5′Cy5-labeled *DHX15* rG4 to screen the compounds from SMM at 50 µM. BRACO19 and DMSO were utilized as positive and negative controls for Z′ factor calculation.

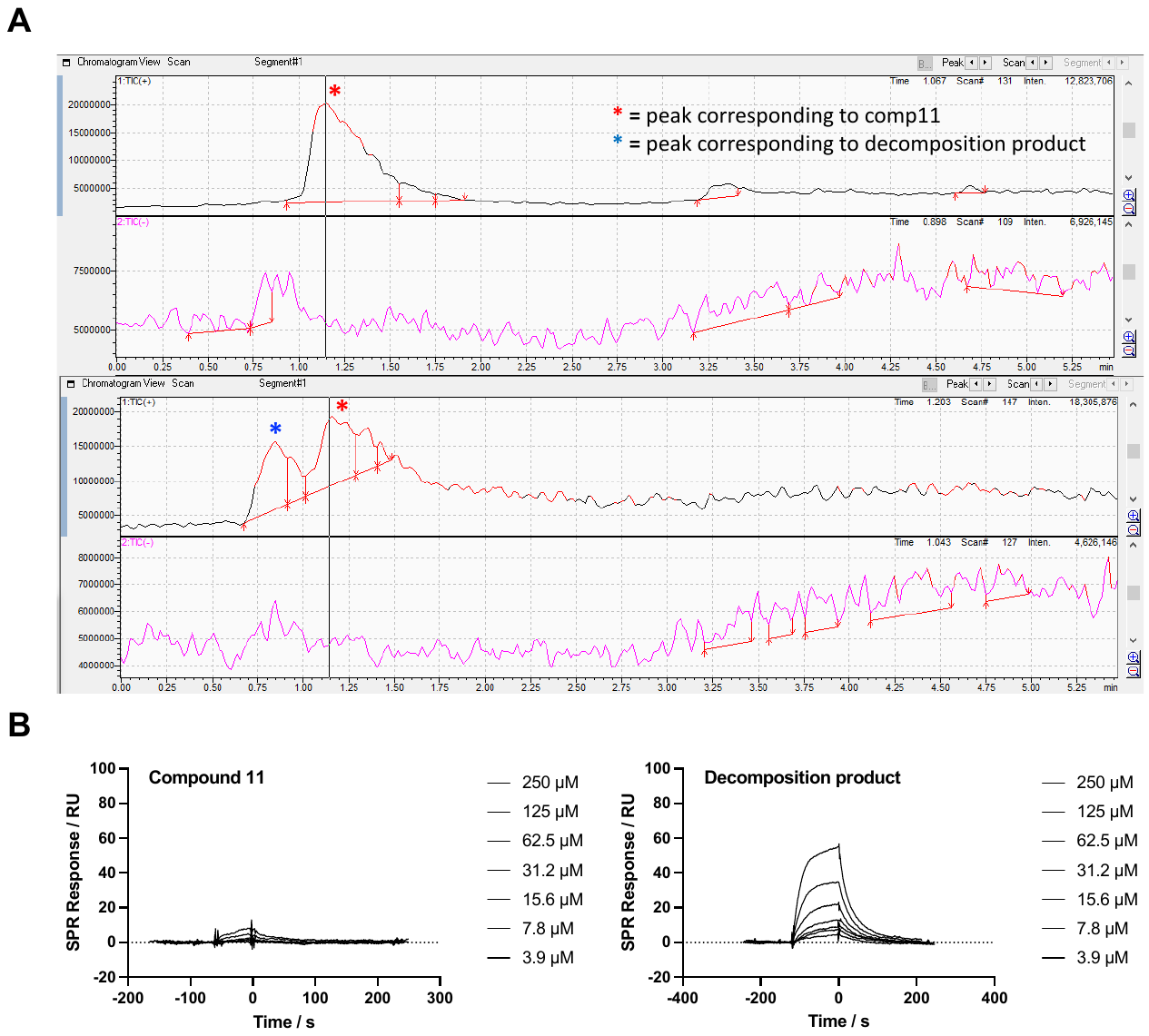

**Supplementary Fig. 4** Evidence for decomposition of compound **11** and binding activity profiling towards *DHX15* rG4. (**A**) UV chromatogram from LC/MS characterization of compound **11** before and after decomposition. (**B**) SPR binding test on compound **11** and the decomposition product.

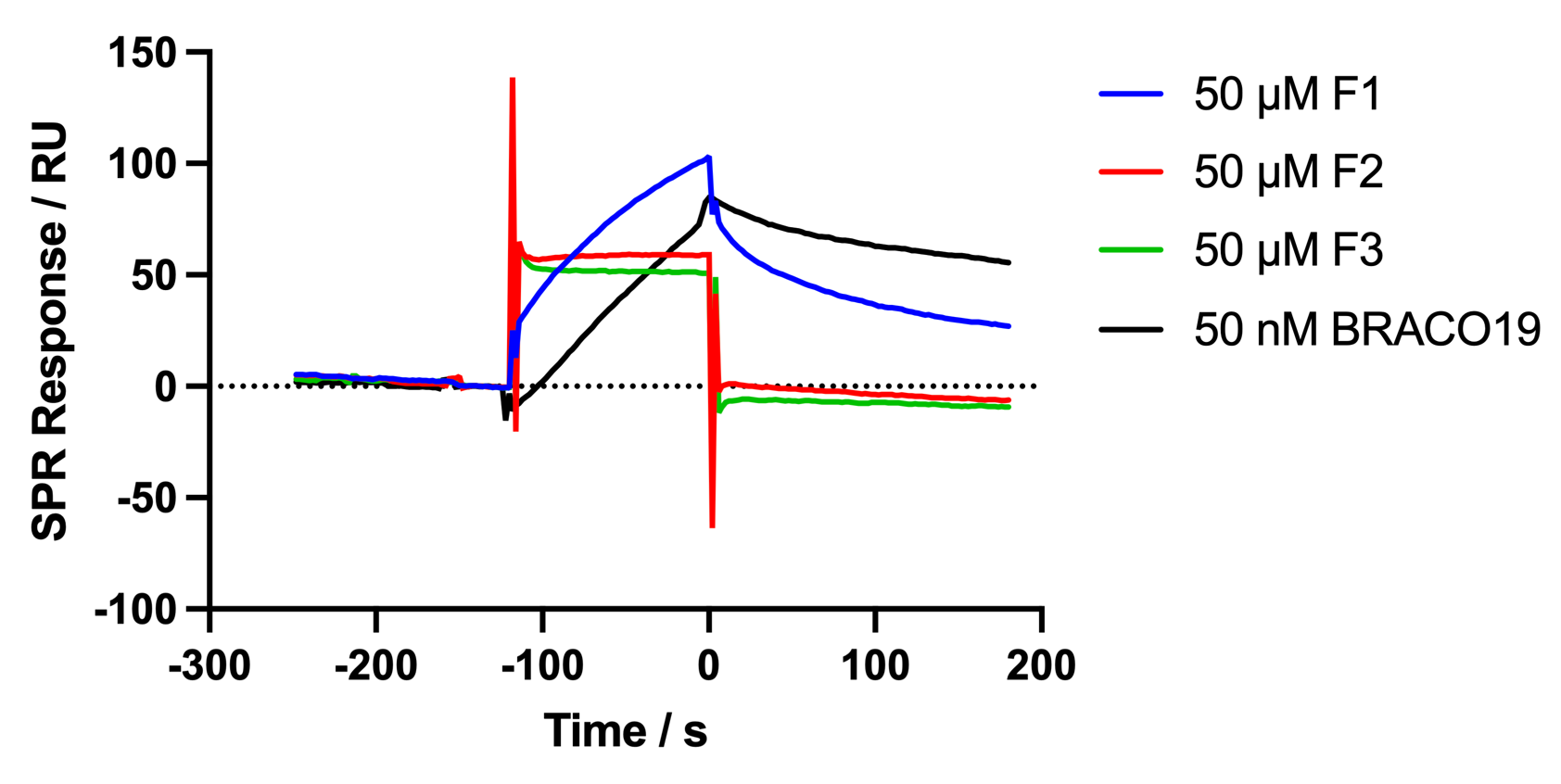

**Supplementary Fig. 5** SPR binding test on the fragments from compound **11**.

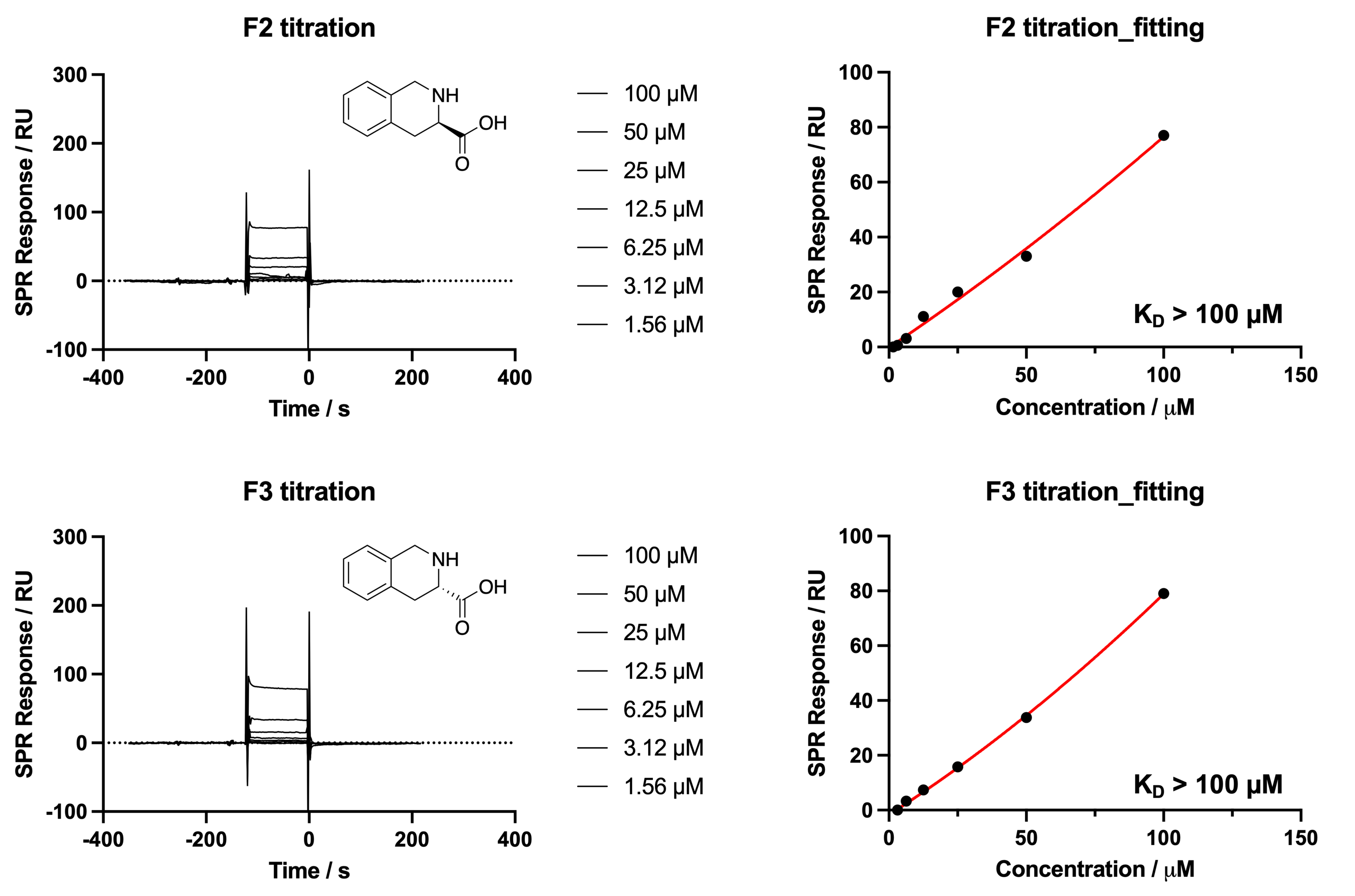

**Supplementary Fig. 6** Titration and K_D_ determination of F2 and F3 by SPR.

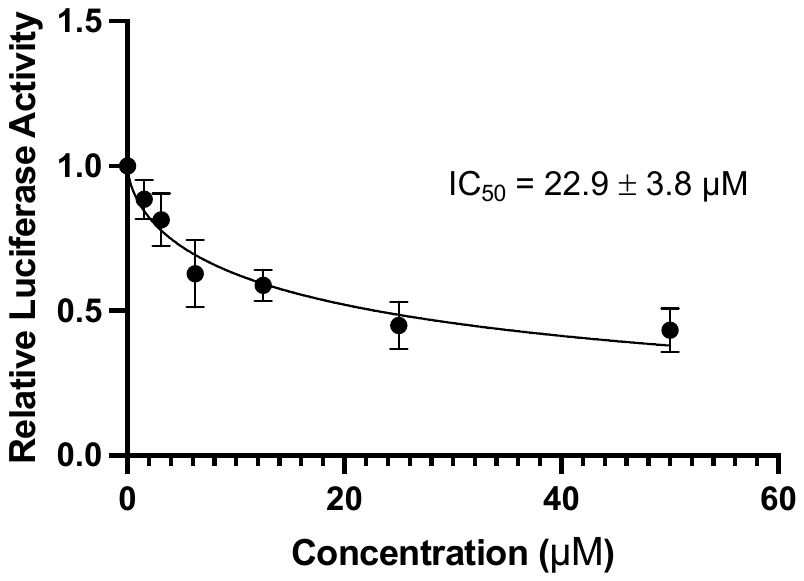

**Supplementary Fig. 7** IC_50_ determination of translation inhibition by F1 in the G4-based luciferase reporter system.

**Chemical Synthesis**

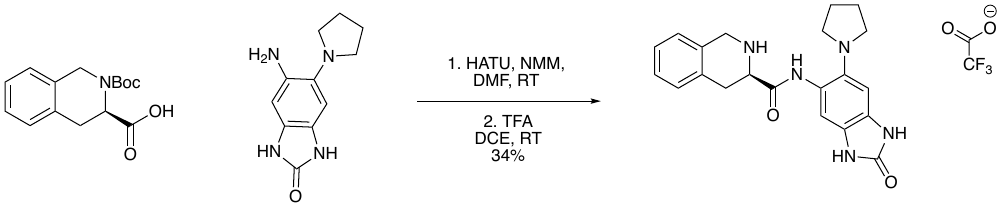

**Compound 11**

To a solution of (*R*)-2-(tert-Butoxycarbonyl)-1,2,3,4-tetrahydroisoquinoline-3-carboxylic acid (200 mg, 0.721 mmol, 1 eq) in dimethylformamide (6 mL) was added HATU (342 mg, 0.901 mmol, 1.25 eq) and N-methylmorpholine (0.16 mL, 1.44 mmol, 2 eq). The mixture stirred at room temperature for 30 minutes, then 5-amino-6-(1-pyrrolidinyl)-1,3-dihydro-2H-benzimidazol-2-one (157 mg, 0.721 mmol, 1 eq) was added. The reaction mixture continued stirring at room temperature for four hours. The reaction was quenched with water (10 mL), then the organic layer was extracted with ethyl acetate (3 x 10 mL). The organic layer and ethyl acetate washings were washed with brine (15 mL) then dried (Na_2_SO_4_). After drying *in vacuo*, the crude product was dissolved in dichloroethane (12 mL). Trifluoroacetic acid (0.83 mL, 10.8 mmol, 15 eq) was added, and the deprotection reaction stirred at room temperature for six hours. The mixture was concentrated down onto celite *in vacuo*. The crude product was purified by reverse phase ISCO flash column chromatography (10-45% acetonitrile + 0.1% TFA in water + 0.1% TFA) to afford 120 mg of white solid (44% yield over two steps). ^1^H NMR (500 MHz, MeOD-*d*_4_): δ 7.37 (s, 1H), 7.34-7.27 (m, 4H), 7.11 (s, 1H), 4.55-4.50 (m, 3H), 3.84-3.81 (m, 4H), 3.62-3.58 (dd, *J* = 4.58, 4.75 Hz, 1H), 3.40-3.33 (m, 1H), 2.30-2.27 (m, 4H); ^13^C NMR (125 MHz, MeOD-*d*_4_): δ 171.1, 158.2, 133.9, 131.7, 131.5, 131.5, 130.0, 129.4, 128.8, 128.7, 127.8, 124.2, 109.4, 103.0, 60.8, 56.9, 45.6, 38.9, 30.4, 25.4; HRMS: (ESI+) m/z calculated for C_21_H_23_N_5_O_2_ [M+H]^+^: 378.1925, found: 378.1936

**NMR Spectra**

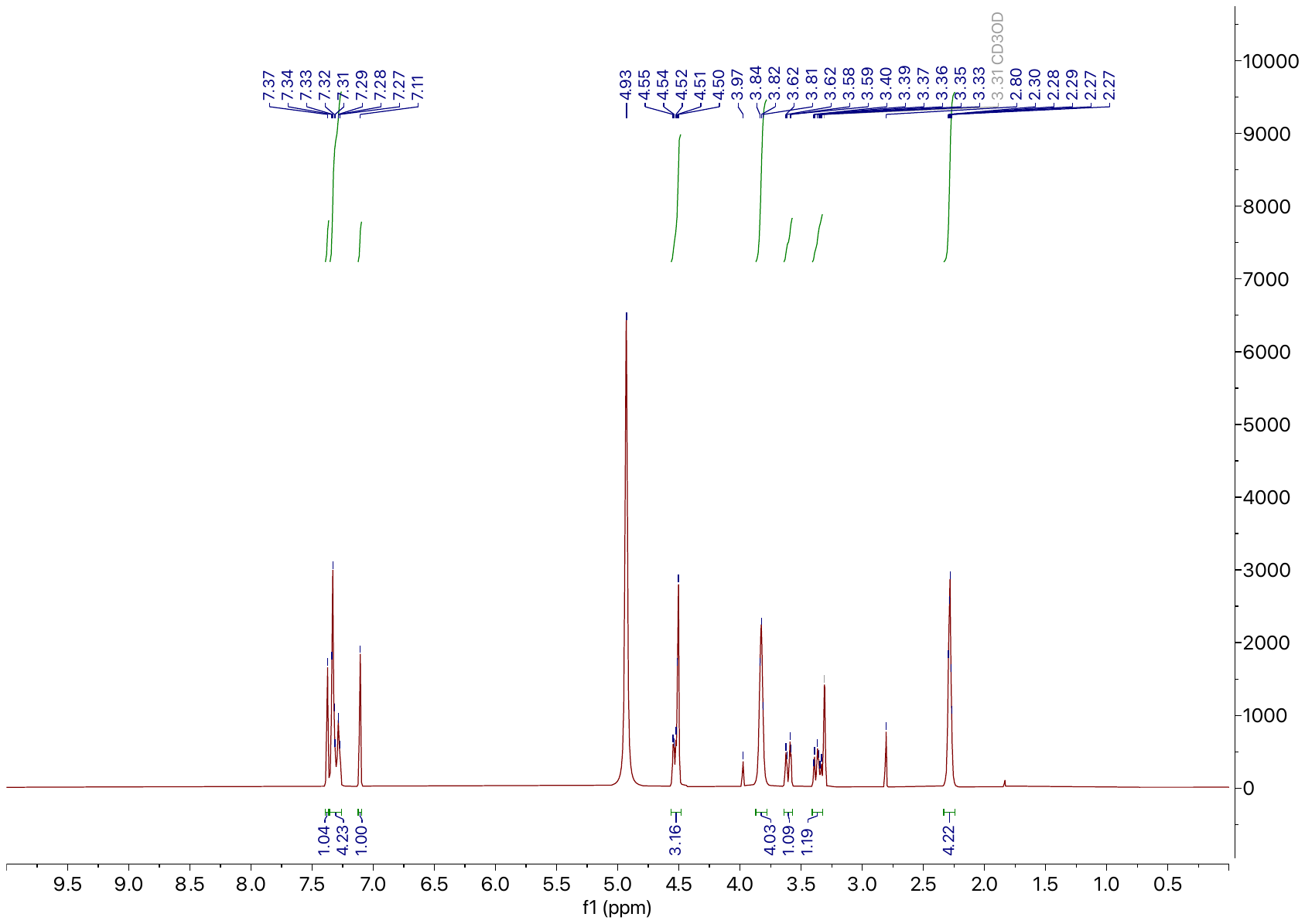

**Compound 11 ^1^H NMR.** 500 MHz, MeOD-*d*_4_

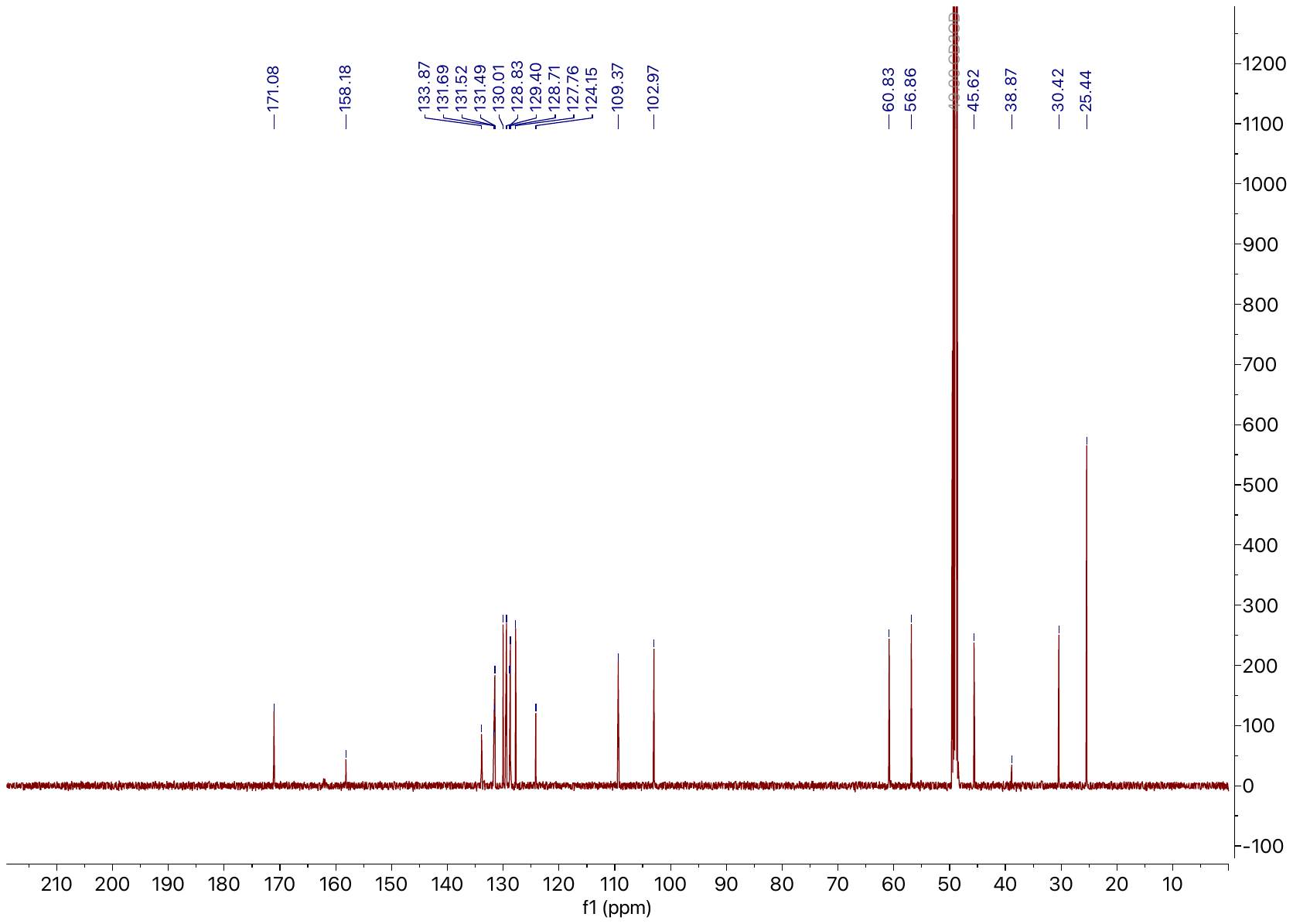

**Compound 11 ^13^C NMR.** 125 MHz, MeOD-*d*_4_
